# Using Ethereum blockchain to store and query pharmacogenomics data via smart contracts

**DOI:** 10.1101/2019.12.16.878488

**Authors:** Gamze Gürsoy, Charlotte M Brannon, Mark Gerstein

**Affiliations:** Program in Computational Biology and Bioinformatics, Yale University, Whitney Avenue, 06520 New Haven, CT, USA; Department of Molecular Biophysics and Biochemistry, Yale University, Whitney Avenue, 06520 New Haven, CT, USA; Department of Computer Science, Yale University, Prospect Street, 06520 New Haven, CT, USA

**Keywords:** blockchain, ethereum, pharmacogenomics data, smart contract, secure storage

## Abstract

**Background:** With the advent of precision medicine, pharmacogenomics data is becoming increasingly critical to patient care. These data describe the relationship between a particular variant in the genome and the response to a drug by the patient. As utilizing this kind of data becomes more integral to medical treatment decisions, appropriate storage and sharing of this data will be critical. A potential way of securely storing and sharing pharmacogenomics data is a smart contract with the Ethereum blockchain. This is an open-source blockchain platform for decentralized applications. A transaction-based, state machine, the “world” of Ethereum maintains user accounts and storage in a network state. Immutable pieces of code called “smart contracts” may be deployed to the Ethereum network and run on the Ethereum Virtual Machine when called by a user or other contract. The 2019 iDASH (Integrating Data for Analysis, Anonymization, and Sharing) competition for Secure Genome Analysis challenged participants to develop time- and space-efficient smart contracts to log and query gene-drug relationship data on the Ethereum blockchain.

**Methods:** We designed a smart contract to store and query pharmacogenomics data (gene-drug interaction data) in Ethereum using an index-based, multi-mapping approach allowing for time and space efficient storage and query. Our solution to the IDASH competition ranked in the top three at a workshop held in Bloomington, IN in October 2019. Although our solution performed well in the challenge, we wanted to improve its scalability and query efficiency. To that end, we developed an alternate “fastQuery” solution that stores pooled rather than raw data, allowing for significantly improved query time for 0-AND queries, and constant query time for 1- and 2-AND queries.

**Results:** We tested the performance of both of our solutions in Truffle (v5.0.31) using datasets ranging from 100 to 1000 entries, and inserting data at 25, 50, 100, and 200 observations at a time. On a private, proof-of-authority test network, our challenge solution requires approximately 70 seconds, 500 MB of memory, and 80 MB of disk space to insert 1000 entries (200 at a time); and 400 ms and 5 MB of memory to query a two-AND query from 1000 entries. This solution exhibits constant memory for insertion and querying, and linear query time. Our alternate fastQuery solution requires approximately 60 seconds, 500 MB of memory, and 80 MB of disk space to insert 1000 entries (200 at a time); and 83 ms and 5 MB of memory to query a two-AND query from 1000 entries. This solution exhibits constant memory for insertion and querying, linear query time for 0-AND queries, and constant query time for 1- and 2-AND queries in a database of up to 1000 entries.

**Conclusion:** In this study we showed that pharmacogenomics data can be stored and queried efficiently on the Ethereum blockchain. Our approach has the potential to be useful for a wide range of datasets in biomedical research; while we focused on gene-drug interaction data, our solution designs could be used to store a range of clinical trial data. Moreover, our solutions could be adapted to store and query data in any field where high-integrity data storage and efficient access is required.

## Background

Pharmacogenomics data, which describes the results of particular gene-drug interactions, allows researchers and physicians to predict how specific patients will respond to a given drug based on the genetic variants they possess. For example, a patient may be more prone to toxic effects from a drug because they possess a variant which limits their ability to clear the drug, and therefore should be prescribed an alternative. The Mayo Clinic likens inclusion of pharmacogenomics data in a patient’s chart to a “flashing, genomic medical alert band,” implying that this data may become as commonplace in basic medical care as wristbands indicating a patient’s allergies or fall risk [1]. Given the growing reliance on this data for medical treatment, its corruption or loss, whether intentional or accidental, has the potential to directly impact medical treatments. It is thus of the essence to develop a robust method for storing, sharing, and updating pharmacogenomics databases in a secure, high-integrity fashion.

The importance of securely storing personal genomic data to protect personal privacy has been widely noted [2]. Pharmacogenomics data presents similar concerns for privacy and immutability. For both genomics and pharmacogenomics, data integrity is critical, as loss, corruption, or alteration of the data would have problematic effects (in the case of genomics, making the wrong diagnosis, and in pharmacogenomics, prescribing the wrong drug). Thus, any method for storing and sharing pharmacogenomics data must prevent it from being lost, changed, or corrupted. A balance between accessibility and privacy is also key to both kinds of data. Researchers and physicians must be able to access genomic data without leaking private information about their patients. They also must be able to store and share gene-drug interaction data during clinical trial phases, while respecting proprietary protocol (i.e. within a pharmaceutical company) or simply sharing only with other groups contributing to the study (i.e. within or between academic institutions). These requirements invite development of creative solutions in order to guarantee secure, robust pharmacogenomics databases.

Blockchain technology is growing in popularity to solve secure data storage problems because of its decentralization, distributed architecture, and immutable linking. Decentralization prevents any single user from controlling the data; distributed architecture eliminates the single point of failure; and immutable linking prevents alteration of past records. Given the requirements for storing pharmacogenomics data discussed above, blockchain technology is an ideal implementation. Some argue that blockchain is a technology fad, and is being used to solve problems that other, simpler technologies could solve, namely a distributed database [3]. However, according to Kuo et al. [“Blockchain distributed ledger technologies for biomedical and healthcare applications,” JAMIA, 2017], blockchain offers features which distributed databases do not, including decentralized management, an immutable audit trail, data provenance, robustness and availability, and security and privacy. These key benefits make blockchain better suited for biomedical applications than other distributed database management systems [4].

A blockchain is a decentralized, distributed, digital ledger comparable to an append-only list linked by cryptographic hashes [5]. The ledger is shared in a “peer-to-peer” network; each node in the network keeps a copy of the list on their computer which syncs to the rest of the network. Nodes in the network submit transactions, which may be verified and added to the chain in a new block. Each block possesses a hash of its contents and the previous block’s hash. Thus, one cannot alter the record, as even the smallest alteration will drastically change the downstream block hashes. The blockchain data structure was first introduced in 2008 by Satoshi Nakamoto (pseudonym) as a digital ledger for transactions of Bitcoin, a now infamous cryptocurrency [5]. However, since this initial application blockchain technology has evolved into a more versatile and dynamic technology, now used for tasks ranging from food distribution tracking to music streaming [6, 7].

Blockchain is increasingly applied to solve real-world problems because of the transparency, immutability, security, privacy, and disintermediation it provides. To achieve transparency, every transaction on the chain is broadcast to all other users in the network, who mine and verify the transactions. Immutability arises because each block contains the hash of all contents in the previous block, preventing any changes to past transactions. Security is achieved through distributed architecture; the transaction history and data stored on the chain is distributed across many nodes in the network, so there is no single point of failure. User privacy is preserved, in a public blockchain network at least, because users can be anonymous, identifiable only by their wallet address. This technology also eliminates the need for a middle man, such as a bank or a music streaming platform, and allows users to transact directly with other users because there is no need for trust in the network [5].

Many of today’s blockchain applications were built to run in Ethereum. Ethereum is a transaction-based state machine which logs modifications to its state in a blockchain. This machine permits development of applications designed for both public and private blockchains through ‘smart contracts’ [8]. Smart contracts are self-executable pieces of code which live in the Ethereum state and trigger transactions when called by a user or another smart contract [9]. Whereas transactions in other blockchain environments, such as bitcoin, were limited in their complexity due to the network state configuration and programming language used, smart contracts are programmed in turing-complete languages such as Solidity, an object-oriented language influenced by JavaScript, C++, Python, and PowerShell [10]. Smart contracts offer transparency (users can verify who deployed the program and ensure they are using the correct version of the program) and immutability (the program cannot be altered and any new versions of the program are accessible to all users). A common example of a smart contract is one which sends currency from one user to another given a certain condition has been met. Yet, there is significant potential to use smart contracts to perform transactions in a variety of contexts and industries.

For the 2019 iDASH competition, we aimed to develop an Ethereum smart contract for storing and querying pharmacogenomics data with time and memory efficiency. While blockchain technology offers several useful features, it is notoriously inefficient and slow when it comes to storing and querying data [5]. Thus, it is challenging to use for everyday applications. We overcome this challenge by using an index-based, multi-mapping data structure in a Solidity smart contract to store the data. Specifically, we store each pharmacogenomics observation in a database mapping in which the key is an index and the value is a struct of the data. We also keep three other indexes, in which the key is a different field (gene name, variant number, and drug name, respectively), and the value is an array holding the indexes that key into the database and match that particular field. This challenge solution allowed for time and memory efficient data insertion and querying by one to three fields. For the sake of scalability, we developed an alternate solution referred to as “fastQuery,” which makes use of a similar query algorithm as our challenge solution, but pools the gene-variant-drug relations in storage during insertion rather than during querying. This fastQuery solution exhibited significantly increased time efficiency for querying by one to three fields.

## Methods

In this study we designed a time/space efficient data structure and algorithms to store and query pharmacogenomics within a smart contract in a private Ethereum blockchain. The data was stored on a small blockchain network with only four nodes. Each data point was inserted to our smart contract as a single transaction. All data had to be stored on-chain. No off-chain data storage was allowed.

### Blockchain technology explained

As outlined earlier, a blockchain is a secure data structure comparable to an append-only list [Fig1A]. Each block in the chain contains a header and a list of valid transactions. The header includes several fields relevant to both the integrity of the data structure, and the parameters of the network. These fields include the block timestamp, block number, mining parameters, and the hash of the previous block’s contents (which links one block to the next). Users or nodes can submit transactions which modify the state of the network, for example by sending value from one user address to another, or storing a value within a smart contract (discussed further below). The network consensus mechanism determines which user will append the transactions to the chain as a new block. The most prominent consensus mechanisms include proof of work, proof of stake, and proof of authority. In a proof of work (PoW) network, nodes “mine” blocks; that is, they find values computationally which, when added to the incoming block’s contents, yields a block hash below that of a set network parameter referred to as the ‘difficulty’. The difficulty is a network-wide parameter which can be varied in order to regulate the rate of block formation. The proof of work mechanism is critical to public cryptocurrency blockchain networks such as Bitcoin because it sets a cost for modifying the blockchain, thereby deterring bad actors from corrupting the chain [5]. Many have criticized the PoW mechanism for being unsustainable in the long run because of the large amounts of energy required to perform computational mining [11]. To address this criticism, the proof of stake mechanism has been proposed [12]. With proof of stake, a network algorithm determines which node will add the block to the chain based on the node’s ‘stake,’ a combination of parameters including their account balance [13]. The PoS mechanism does not require heavy computation, thereby reducing the energy usage in the network. These consensus mechanisms are essential for a public blockchain network where anyone in the world can run a node and potentially modify the chain. However, in a private, permissioned network, these mechanisms can be replaced with a simple proof of authority mechanism [5]. PoA is a modified version of PoS with identity as the only ‘stake’ [13]. We tested our smart contracts on a private test network using proof of authority.

**Fig. 1:**
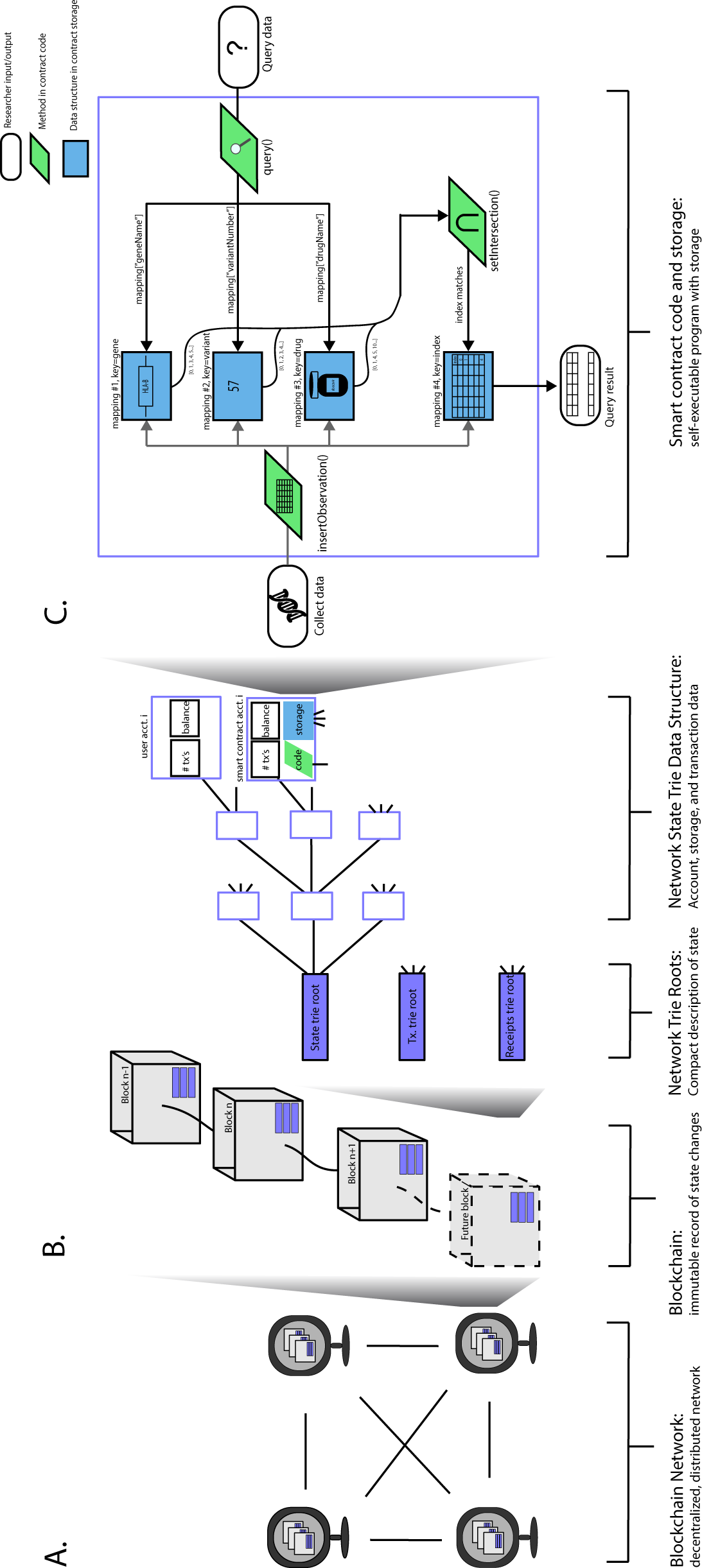
Ethereum blockchain and smart contracts. (a) A blockchain network consists of a decentralized, distributed digital ledger shared in a peer-to-peer fashion. (b) A blockchain can be visualized as a string of blocks linked by cryptographic hashes. Blocks contain data about the state of the network stored in a trie root. The state trie data structure in Ethereum stores information for user and smart contract accounts. (c) Flowchart of the insertion and query processes in our challenge solution smart contract.

### Ethereum blockchain and smart contracts

A broad view of Ethereum might divide the world into three parts: blockchain, network trie roots, and trie data structures [Fig1B]. The blockchain logs the network’s state at specific times, after transactions have altered the state. The state of the network is stored in merkle patricia trees each of which possess a top hash. Blocks in the chain store these top hash values, but do not store all the data in the blockchain network (it would be far too large). The state data is stored in a database layer using leveldb [14, 15]. User account data and smart contract data (including the code and the actual data inserted via the code) are stored in these trie data structures, which are synced by “full” and “archival” nodes only (nodes which require significantly more computational power and storage) [15]. These nodes are integral to the health of the network.

Ethereum can handle a wide range of transactions via smart contracts, self-executable turing-complete programs which run in the Ethereum Virtual Machine (EVM) and maintain state in their own storage [8]. The EVM has a stack-based architecture, and can either store things on the stack (e.g. byte-code operations), in memory (e.g. temporary variables within functions), or in storage (e.g. permanent variables holding database entries). Each smart contract can read and write to its own storage only. In order to discourage developers from writing inefficient or unwieldy smart contracts, there is a ‘gas’ cost associated with each storage and retrieval command. Just as blockchain users have an address, a smart contract’s state resides at a particular unique address in the global state of the Ethereum network, which users can call. If a user does not have enough gas, the contract call and corresponding transaction cannot be completed. Smart contracts provide an opportunity to develop applications with complex functionality in a blockchain network. We leveraged the flexibility of smart contract programming to create a challenge solution and alternate fastQuery solution that insert pharmacogenomics data in a custom way in contract storage to maximize storage and query efficiency.

### Network Configuration

We developed and tested our solutions in Truffle v5.0.31, a command line tool for Ethereum, which provided us with a built-in JavaScript test environment. Within this setup, we tested insertion and querying with up to 1000 entries in the database.

### Chain initialization

Chain initialization in Truffle is automated, and only requires setting basic parameters such as gas limit and network name in a config file. We used the default gas limit and price values for the development network (4712388 and 100 gwei, respectively).

### Challenge solution: an index-based, multi-mapping smart contract

Our challenge solution utilizes four core storage mappings, linked to one another by an index assigned to each inserted observation in the database [Fig1C]. Mappings are similar to hash tables, and allow efficient key-value lookup. In the first of the four mappings, which we call the database, we store the pharmacogenomics data and assign to each entry a unique index to serve as the mapping key. We store each observation in its own struct, a composite data type which can hold multiple fields. In the three other mappings, we use the gene names, variant numbers, and drug names as keys to an array of relevant indexes. Thus, given a gene name key, one can retrieve a list of indexes that could key into the database mapping and return an observation matching that particular gene name. We chose this implementation in order to reduce the number of loops required to check the data in the database and return matches, and thereby achieve time/memory efficient querying.

### Challenge solution-insertion

Data can be inserted into a smart contract via one-line commands in a JavaScript console or from an external script. We wrote an insertion function within the contract to insert a single observation for a given gene name-variant number-drug name combination. In our case the observation consists of the gene name, variant number, drug name, outcome (improved, unchanged, or deteriorated), suspected gene-outcome relation (true/false), and serious side effect (true/false). Upon passing in the observation, the function executes the following steps, (1) Convert each field of the observation to the desired data type for storage (e.g. it is more efficient to store strings as bytes32 types in storage); (2) if the gene name, variant, drug combination does not already exist in the database, add it to an array holding only the unique gene name, variant, and drug name combinations; (3) Use the gene name, variant number, and drug name to key into their respective mapping and append to the value array the counter as the index, where counter is a globally updated variable; (4) push to the database mapping a struct with key-value pair: counter-[struct holding the observation]; (5) increment counter variable by one.

We store the gene name, variant number, and outcome fields as bytes32 variables, a fixed-size bytes array that uses less ‘gas’ in Ethereum relative to strings and is compatible with basic utilities in Solidity (for example, it is simpler to check the length of a bytes variable than a string variable using Solidity). We store the drug name as a string because many drug names are lengthy and can exceed the 32-character limit of the bytes32 data type. Booleans are straightforward in Solidity, and addresses are types to conveniently store the user or contract addresses on the blockchain.

### Challenge solution-query algorithm

Our challenge solution can handle both three field queries, where the fields are gene name, variant number, drug name, and wildcard queries. One can query by any combination of these three fields, or simply specify gene=“*”, variant= “*”, drug= “*”, which should return the entire contents of the database. To query, we first check how many fields have been queried, and for those that were, we use the query fields to key into the appropriate mapping and return the value associated with that key– an array of integer indexes corresponding to the index of the data in the database mapping. If all fields were “*” we use all indexes currently in use to return the data from the database mapping. Otherwise, we determine which of the index-arrays has the minimum length, and loop through it (since its contents will limit the results of the set intersection). For each entry in this minimum-length index array, for every other field that was searched we loop through that index array and check whether it matches the index in the outer loop. At the end of the outer loop, if the number of matches is equal to the number of fields searched, then this integer is used to key into the database mapping and grab the struct value, which is then saved to a memory array. We then loop through the array of unique combinations and for each one pool the structs in the search results that match that unique combination. This allows us to output data in a useful way: for a given gene name, variant number, and drug name, how many observations are there, what number and percentage of cases saw serious side effects, what number and percentage suspected an interaction between the gene and drug, and what number and percentage saw an improved, deteriorated or unchanged outcome when administered the drug.

### fastQuery solution: a pooled, index-based, multi-mapping smart contract

Rather than storing each observation in its own struct, our alternate, fastQuery solution stores a single struct for each unique gene-variant-drug relation. We utilize four core storage mappings linked to one another by an index assigned to each inserted observation in the database, and an additional fifth mapping for linking indexes to their unique gene-variant-drug relation. This alternate design reduces the number of indexed entries in the database, and prevents indefinite database growth (there is an infinite number of raw observations that can be inserted into our challenge solution, but a finite number of unique gene-variant-drug relations that exist). We chose this implementation in order to reduce the length of loops required to check the data in the database and return matches, and thereby further improve the query time.

### fastQuery solution-insertion

We wrote a new insertion function to insert a single observation for a given gene name-variant number-drug name combination. Upon passing in the observation, the function executes the following steps, (1) Convert each field of the observation to the desired data type for storage; (2) Use the input gene, variant, and drug as a combination key into the fifth mapping, which returns the index for that unique relation; (3) Use the index as a key into the database mapping and check whether this relation already exists in the database; (4) if not, fill in the name information from the input data, add the gene, variant, and drug names as keys in their respective mappings with the index as the value, and increment the global index counter by 1; (5) increment the total count, improved count, deteriorated count, suspected relation count, and side effect count fields of the relation struct in the database based on the inserted data. We store the variables as the same data types used in our challenge solution.

### fastQuery solution- query algorithm

Like our challenge solution, our fastQuery solution can handle both three field queries, where the fields are gene name, variant number, drug name, and wildcard queries. The first part of our query algorithm remains very similar to the challenge solution; we first check how many fields have been queried, and for those that were, we use the query fields to key into the appropriate mapping and return the value associated with that key– an array of integer indexes corresponding to the index of the data in the database mapping. If all fields were “*” we use all indexes currently in use to return the data from the database mapping. Otherwise, we determine which of the index-arrays has the minimum length, and loop through it (since its contents will limit the results of the set intersection). For each entry in this minimum-length index array, for every other field that was searched we loop through that index array and check whether it matches the index in the outer loop. At the end of the outer loop, if the number of matches is equal to the number of fields searched, then this integer is used to key into the database mapping and grab the struct value, which is then saved to a memory array. At this point, our query algorithm differs from the challenge solution. In fastQuery, we do not need to loop through an array of unique gene-variant-drug combinations because the data are already stored pooled in these unique relations. Thus, we simply loop through the matches, convert the count data to percentages, and output the search result.

## Results

We present two proof-of-concept solutions for storing pharmacogenomics data observations of up to 1,000 entries: our challenge solution, and an alternate fastQuery solution with improved query times. Both solutions were measured for its accuracy, time and space efficiency, and scalability [Fig. 2 and Fig. 3].

**Fig. 2:**
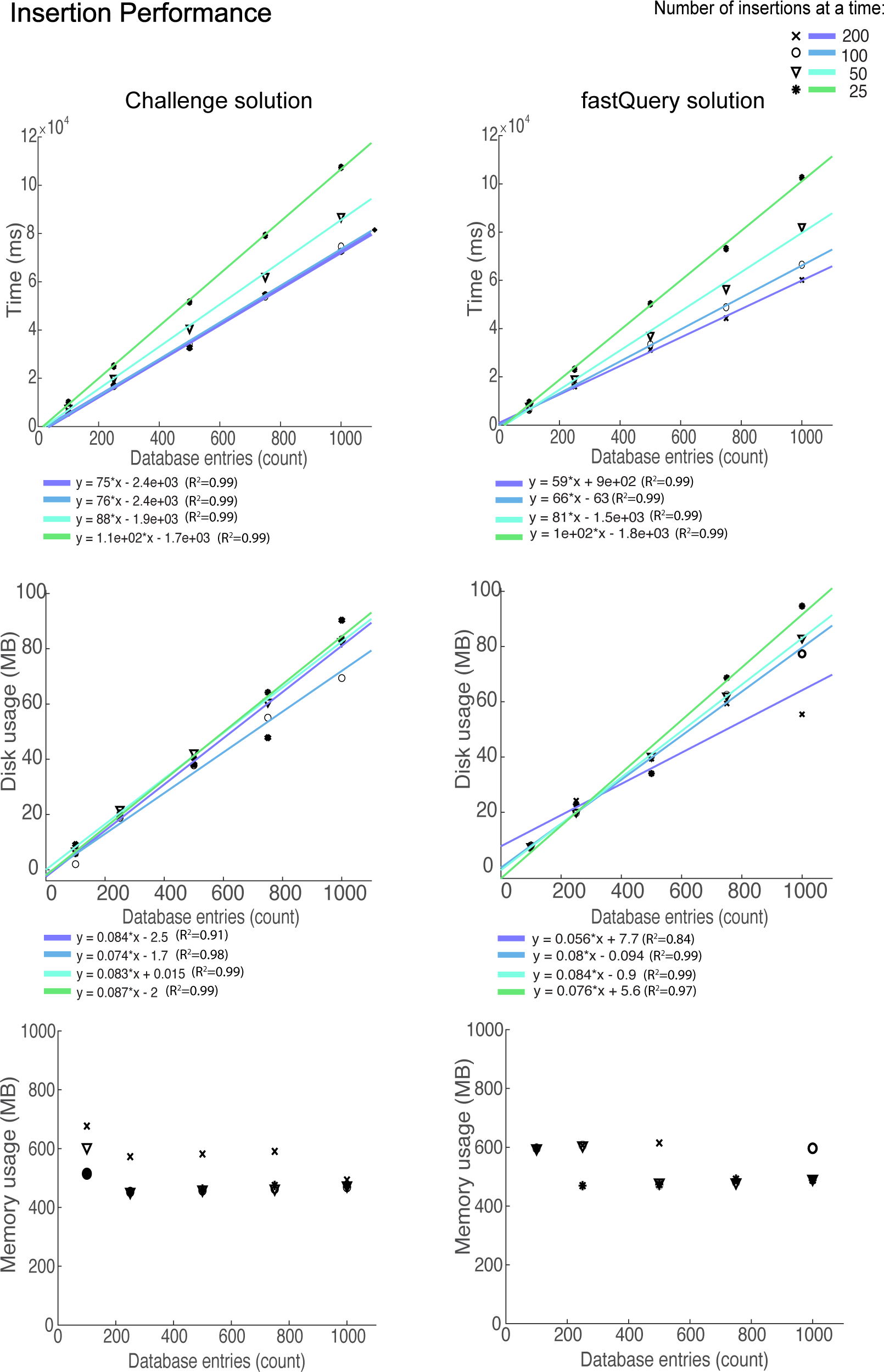
Insertion results. Time, memory, and disk usage of inserting data into our two smart contract solutions.

**Fig. 3:**
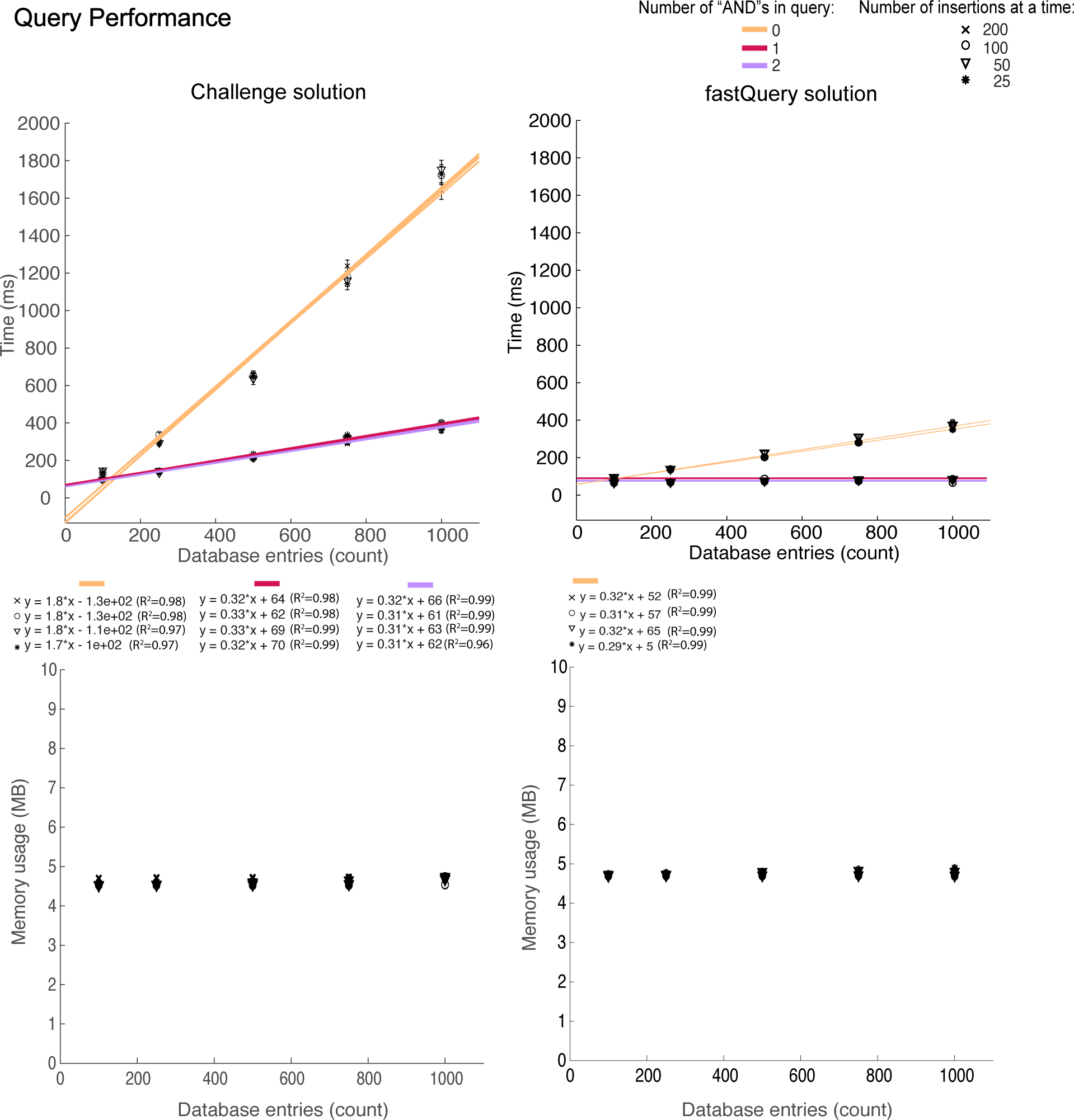
Query results. Time, memory, and disk usage of querying data from our two smart contract solutions. Comparison of query performance for zero-, one-, and two- AND queries.

### Accuracy

The Truffle environment allows testing from custom JavaScript scripts. Using assertions in JavaScript, we checked that the query results matched the fields queried for 100 random queries with zero, one, and two “AND”s. the challenge and fastQuery solutions.

### Time and space efficiency and scalability

#### Insertion

We measured the time, memory, and disk usage required to insert increasing amounts of data into contract storage, where each insertion is a single empirical observation of a gene-variant-drug interaction (for example, gene=”CYP3A5”, variant=52, drug=pegloticase, outcome=UNCHANGED, suspected gene-drug-relation=true, serious side-effect = true) [Fig. 2]. We measured insertion time when inserting 25, 50, 100, and 200 entries at a time (with 1 s pauses between batches, subtracted from the reported times). We found that our challenge solution takes approximately 67 s to insert 1000 observations, regardless of the number of insertions at a time, with a linear time complexity. We found that the memory requirement for inserting 1,000 entries into the chain is around 500 MB regardless of the number of insertions at a time, which remains constant with increasing number of entries. We found that disk space usage increases linearly with database entries, with approximately 80MB required to store 1000 entries.

The fastQuery solution takes approximately 56 s regardless of the number of insertions at a time, with a linear time complexity [Fig. 2]. The memory requirement for inserting 1,000 entries into the chain is around 500 MB regardless of the number of insertions at a time, which remains constant with increasing number of entries. Disk space usage increases linearly with database entries, with approximately 80MB required to store 1000 entries. Thus, fastQuery shows improvement for insertion time, but performs the same as the challenge solution for memory and disk usage.

#### Querying

We measured the performance of our query algorithm by testing the time and memory required to do a three-field query in a database with an increasing number of entries for 100 random queries [Fig 3]. We found that our challenge solution takes an average of approximately 300 ms to complete a two-AND query from a database of 1000 entries with an estimated linear time complexity. We also measured the memory requirement for the same query, and found that it was approximately 5 MB per query and constant with increasing database entries.

Our fastQuery algorithm showed improved query time [Fig 3]. We found that it takes approximately 83 ms to complete a two-AND query from a database of 1000 entries, and constant time with increasing number of database entries. It showed similar performance to the challenge solution for query memory requirement (approximately 5MB per query and constant with increasing database entries.

#### Effect of varying “ANDs” in query

We investigated whether the time and memory efficiency of our two solutions varied with the number of “AND”s in a query [Fig 3]. We checked whether for a database of 1000 entries, changing the number of fields queried affects the time and memory requirement. We found that for one- and two-AND queries in the challenge solution, it takes an average of approximately 200 ms to query from a 1000-entry database, while a 0-AND query takes an average of approximately 1.6 s to do the same. We found memory usage to be constant for up to 1000 entries, and approximately 5MB per query regardless of the number of ANDs in the query. For the fastQuery solution, we found that for one- and two-AND queries take an average of approximately 85 ms to query from a 1000-entry database, while a 0-AND query takes an average of approximately 400 ms to do the same [Fig 3]. As with the challenge solution, we found memory usage to be constant for up to 1000 entries, and approximately 5MB per query regardless of the number of ANDs in the query.

## Discussion

High-integrity, secure data maintenance is a major concern in biomedical research. In the case of pharmacogenomics, the data collected from clinical trials directly impacts medical treatment decisions. The integrity and security of the data is therefore critical, as loss or corruption will lead to misguided medical care. The 2019 IDASH Secure Genome Analysis competition proposed using smart contracts in a private Ethereum blockchain [16]. Such a solution would protect the data from loss in a single point of failure scenario and from accidental or intentional corruption. It also has broader implications, showing the potential for applying blockchain technology to solve real-world problems beyond cryptocurrency.

In this study, we presented two proof-of-concept solution to store pharmacogenomics data using the Ethereum blockchain platform. Both solutions addresses the need for security and accessibility in sharing these data, but also the practical need for time and memory efficiency for use in the real world. We showed that blockchain technology can not only offer security and immutability, but also efficiency and practicality. Although we were able to develop two efficient solutions as Ethereum smart contracts, development in Ethereum is far from easy. Setting up a private blockchain in Ethereum requires expert knowledge in the platform, and deploying the contract is a complex process. This process can be condensed into an external script, which reduces the need for expertise to use the platform. However there are still issues with bugs in Ethereum software such as in web3.js, the JavaScript API for Ethereum. We were able to overcome these software issues, but acknowledge the need for more stability in this platform before researchers begin using it for a shared database.

In summary, we presented a challenge solution for storing and querying pharmacogenomics data on the Ethereum blockchain; and an alternate fastQuery solution with significantly improved query time. Our solutions demonstrate the potential for blockchain technology in the medical research community, but could be applied to a variety of other store and query problems.

## Conclusion

In conclusion, we showed that pharmacogenomics data can be stored and queried efficiently using an Ethereum blockchain smart contract.

## Ethics approval and consent to participate

Not Applicable.

## Consent for publication

Not applicable.

## Availability of data and material

Source codes for insertion and querying, example input file, queries used in this study and the results for insertion and querying can be accessed at https://github.com/gersteinlab/iDASH19bc under GPL v3.0 license.

## Funding

This work is supported by US National Institutes of Health U01 EB023686 grant. Publication costs were funded by US National Institutes of Health U01 EB023686 grant.

## Competing interests

The authors declare that they have no competing interests.

## Author’s contributions

G.G. and M.G designed the project and led the overall analysis. G.G and C.M.B designed the solutions and wrote the code. G.G, C.M.B and M.G. analyzed the results. G.G, C.M.B and M.G. wrote the manuscript. All authors read and approved of the final manuscript.

## Acknowledgements

We thank iDASH organizers for providing an avenue to develop secure genome analysis applications.

## References

1. Mayo Clinic Center for Individualized Medicine. Pharmacogenomics: Drug-Gene Alerts https://www.mayo.edu/research/centers-programs/center-individualized-medicine/patient-care/pharmacogenomics/drug-gene-alerts.

2. Erlich, Y., Narayanan, A.. Routes for breaching and protecting genetic privacy. Nature Reviews Genetics, 2014;15;409–421.

3. Kharpal, A. Blockchain is ‘one of the most overhyped technologies ever,’ Nouriel Roubini says. https://www.cnbc.com/2018/03/06/blockchain-nouriel-roubini-one-of-the-most-overhyped-technologies-ever.html. 2018.

4. Kuo, T., Kim, H., Ohno-Machado, L. Blockchain distributed ledger technologies for biomedical and health care applications. JAMIA, 2017;24(6):1211–1220.

5. Nakamoto S. Bitcoin: A Peer-to-Peer Electronic Cash System. bitcoin.org/bitcoin.pdf, 2008.

6. Browne, R. IBM partners with Nestle, Unilever and other food giants to trace food contamination with blockchain. https://www.cnbc.com/2017/08/22/ibm-nestle-unilever-walmart-blockchain-food-contamination.html, 2017.

7. ConsenSys Ujo and Capitol Records bring blockchain innovation to music. https://media.consensys.net/consensys-ujo-and-capitol-records-bring-blockchain-innovation-to-music-319f2c649790, 2018.

8. Wood, G. Ethereum: a secure decentralised generalised transaction ledger byzantium version https://ethereum.github.io/yellowpaper/paper.pdf, 2019.

9. A next-generation smart contract and decentralized application platform https://github.com/ethereum/wiki/wiki/White-Paper, 2019.

10. Solidity https://solidity.readthedocs.io/en/v0.5.12/, 2019.

11. Zochowski, M Why proof-of-work is not viable in the long-term https://medium.com/logos-network/why-proof-of-work-is-not-viable-in-the-long-term-dd96d2775e99, 2019.

12. King, S., Nadal, S. PPCoin: Peer-to-peer crypto-currency with proof-of-stake https://decred.org/research/king2012.pdf, 2012.

13. POA Network Proof of authority: consensus model with identity at stake. https://medium.com/poa-network/proof-of-authority-consensus-model-with-identity-at-stake-d5bd15463256, 2017.

14. McCallum, T. Diving into Ethereum’s world state. https://medium.com/cybermiles/diving-into-ethereums-world-state-c893102030ed, 2018.

15. The Ethereum world – how its data is stored https://github.com/tpmccallum/ethereumdatabaseresearchandtesting/blob/master/leveldb 2018.

16. IDASH Privacy and Security Workshop 2019 – secure genome analysis competition http://www.humangenomeprivacy.org/2019/competition-tasks.html, 2019.

